# Novel syngeneic animal model of tobacco-associated oral cancer reveals the activity of *in situ* anti-CTLA-4

**DOI:** 10.1101/672527

**Authors:** Zhiyong Wang, Victoria H. Wu, Michael M. Allevato, Mara Gilardi, Yudou He, Juan Luis Callejas-Valera, Lynn Vitale-Cross, Daniel Martin, Panomwat Amornphimoltham, James Mcdermott, Yusuke Goto, Alfredo A. Molinolo, Andrew B. Sharabi, Ezra E. W. Cohen, Qianming Chen, J. Guy Lyons, Ludmil B. Alexandrov, J. Silvio Gutkind

## Abstract

Head and neck squamous cell carcinoma (HNSCC) is the sixth most common cancer worldwide. Tobacco use is the main risk factor for HNSCC, and tobacco-associated HNSCCs have poor prognosis and response to available treatments. Recently approved anti-PD-1 immune checkpoint inhibitors showed limited activity (≤20%) in HNSCC, highlighting the need to identify new therapeutic options. For this, mouse models that accurately reflect the complexity of the HNSCC mutational landscape and tumor immune environment are urgently needed. Here, we report the first mouse HNSCC model system that recapitulates the human tobacco-related HNSCC mutanome, in which tumors grow when implanted in the tongue of immunocompetent mice. These HNSCC lesions have similar immune infiltration and response rates to anti-PD-1 (≤20%) immunotherapy as human HNSCCs. Remarkably, we found that >70% of HNSCC lesions respond to intratumoral anti-CTLA-4. This syngeneic HNSCC mouse model provides a platform for the development of novel immunotherapeutic options for HNSCC.

## Introduction

Tobacco smoking claims the lives of more than 6 million people every year worldwide and is the leading cause of cancer deaths in the U.S^1,2^. Tobacco use has been associated with at least 17 types of cancer, primarily in the lung, as well as with carcinomas arising in the oral cavity, pharynx, and larynx, often referred to as squamous cell carcinomas of the head and neck (HNSCC)^1,2^. HNSCC is a significant public health issue, with more than 65,400 new cases resulting in 14,600 deaths in 2019 in the U.S. alone^3^. The main risk factors include tobacco use and human papillomavirus (HPV) infection, the latter of which is predicted to diminish in the future due to successful vaccination campaigns^4,5^. Depending on the stage of disease, HNSCC is typically treated with surgery, radiotherapy, chemotherapy, or a combination of these interventions. These standard therapies result in a five-year survival of approximately 63%, but patients with more advanced stages have higher rates of mortality (5 year survival <50%) and require multimodality treatments, which can lead to occurrence of significant long-term side effects and lower quality of life^6,7^. Despite this more aggressive regimen, up to 30-60% of HNSCC patients develop tumor recurrence, and often succumb to the disease^8^. The recent elucidation of the genomic alterations underlying HNSCC progression and new immunotherapeutic strategies may provide an opportunity for the development of more effective treatment options for HNSCC.

Revolutionary breakthrough discoveries in cancer immunology have demonstrated that a patient’s own immune cells can be manipulated to target, attack, and destroy cancer cells^9-11^. A key emerging mechanism of tumor immune evasion involves T cell exhaustion, whereby T cell reactivity is impaired due to activation of T cell checkpoints, including PD-1 by its ligand, PD-L1 that is expressed by macrophages and some cancer cells, including HNSCC, restraining T cell activation (reviewed in^12^). Indeed, immune checkpoint blockade (ICB) by new immunotherapeutic agents such as pembrolizumab and nivolumab (anti-PD-1) have recently demonstrated potent anti-tumor activity in a subset of HNSCC patients^13-16^. However, one-year survival and response rates of anti-PD-1 in HNSCC were only 36% and 14%, respectively, which highlights the urgent need to identify novel therapeutic options to increase the effectiveness of ICB for the >80% of patients that do not have an objective response to anti-PD-1/PD-L1 treatment^13-15^.

Animal models with a full functioning immune system that also properly resemble human HNSCC etiology and mutational landscape are desperately needed to accurately recapitulate the complexity of the tumor immune microenvironment (TIME), thereby accelerating the search for new immune therapeutic options. Here, we report the first syngeneic murine HNSCC cell panel that recapitulates typical human tobacco-related HNSCC genomic alterations and mutational landscape, and we show that these cells form squamous carcinomas (HNSCC) when implanted orthotopically in the tongue of immune competent C57Bl/6 mice. These HNSCC lesions have immune infiltration and response rates to anti-PD-1 therapies (≤20%) similar to those of human HNSCCs, thereby providing a platform for the evaluation of new immune oncology (IO) options for HNSCC treatment.

## Results

### Novel syngeneic HNSCC animal models exhibiting tobacco carcinogen-related mutational signatures and genomic landscape

Tobacco smoke contains a number of harmful carcinogens that drive tumorigenesis, the exposure to which strongly correlates with cancer incidence^17^.While tobacco-associated cancers are generally characterized by high mutation frequencies^18^, we have recently reported that they can be defined by very specific set of mutational signatures^19^. We have also described the optimization of a carcinogen-induced oral cancer mouse model in which the compound 4-nitroquinoline-1 oxide (4NQO), a DNA adduct-forming agent that causes DNA damage and promotes oral cancer initiation and progression^20^. This model has been used extensively to study HNSCC progression and preventive and treatment therapeutic options^21-23^. However, its direct relevance to human HNSCC has not been previously established. To begin developing syngeneic HNSCC animal models, we first isolated 4 representatives murine HNSCC cell lines from primary 4NQO-induced tumors in the tongue of C57Bl/6 mice (designated 4MOSC1-4, short for 4NQO-induced Murine Oral Squamous Cells) (**Fig. 1a**). The use of SigProfiler^24,25^ to analyze exome DNAseq of these HNSCC cells revealed a remarkable 93.9 % similarity with human cancer signature 4, which is strictly associated with tobacco smoking, including in HNSCC, esophageal cancer, and lung cancer^19^ (Pearson correlation > 0.93) (**Fig. 1b** and individual 4MOSC cells in **Supplementarly Fig 1**). This similarity between 4NQO-induced mutational patterns and tobacco extended to the presence of a transcriptional strand bias (**Fig. 1c**), which reflects rate of substitution type on each nucleotide. In contrast, the mutational signature of SCC caused by DMBA, a carcinogen found in tobacco smoke that is the most widely used agent for experimental carcinogenesis studies^26^, showed only 39.7% similarity with human cancer signature 4. This suggests that 4NQO-induced SCC lesions better reflect human tobacco-related human HNSCC. Indeed, these cells also exhibit typical HNSCC histology and mutations impacting *Trp53, Fat1-4, Keap1, Notch1-3, Kmt2b-d*, and others, which represent some of the most frequently altered gene pathways in HPV-human HNSCC (**Fig. 1d-e**, and **Supplementary Table 1**). Of note, similar to HPV(-) HNSCC samples from TCGA, all four 4MOSC cells exhibit typical inactivating *Trp53* mutations in its core DNA binding domain, including hot spot residues (G245, and R248) that result in loss of tumor-suppression and gain of tumorigenesis and invasiveness^27^.

**Fig. 1.**
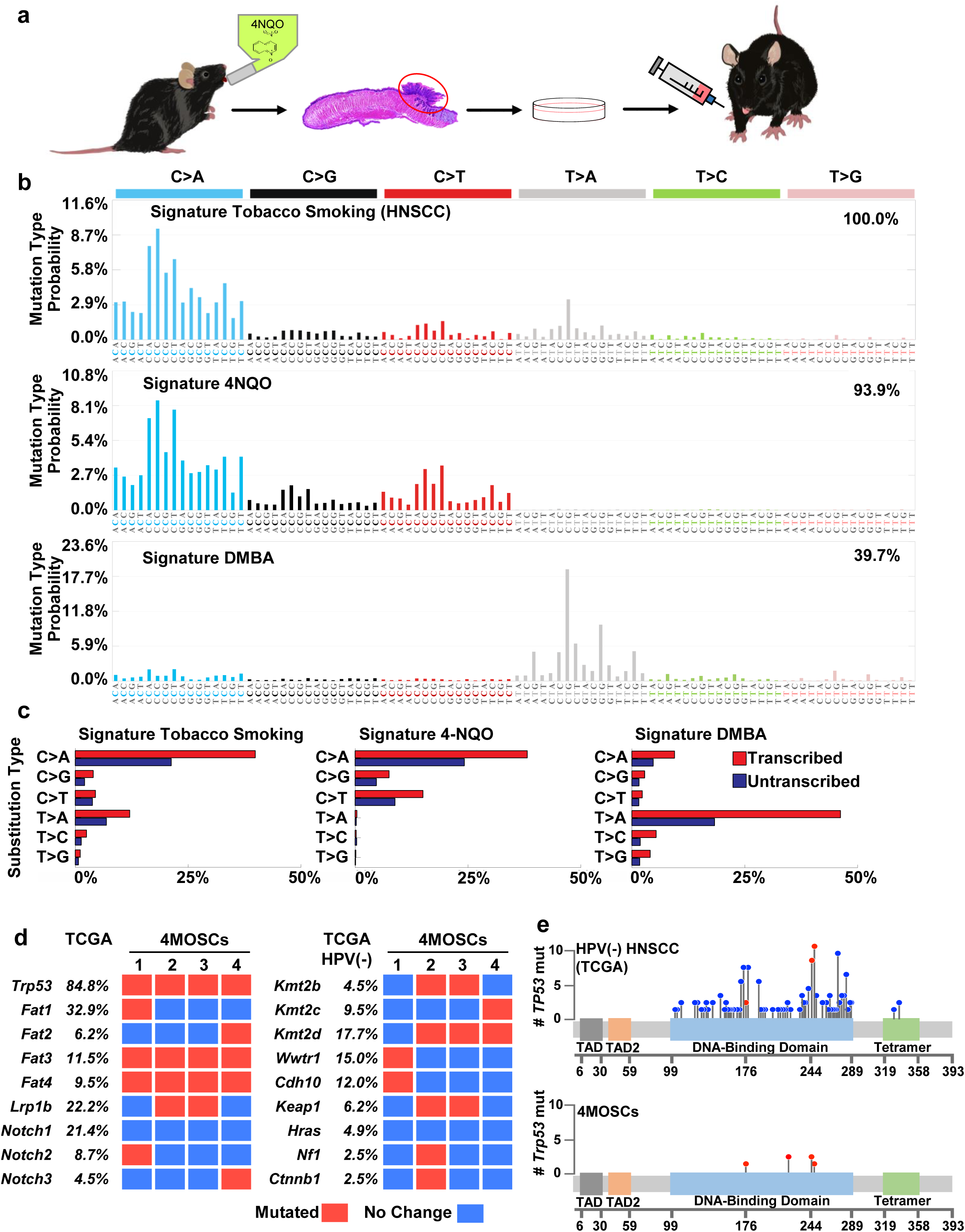
Development of a novel syngeneic mouse model for oral squamous cell carcinoma. **(a)** Experimental scheme of 4NQO syngeneic model. C57Bl/6 Mice were given 4NQO (50 μg/mL) in the drinking water for 16 weeks and then regular water until week 22. Cells were isolated from the lesions, cultured, and then implanted into the tongue of wild-type C57Bl/6 mice. **(b)** Mutational signatures associated with tobacco smoking. The somatic mutational profiles of the four lesions from mice exposed to 4NQO were correlated to known mutational signatures in human cancer (Pearson correlation > 0.93)^19,24^. Top, Signature 4 extracted from cancers associated with tobacco smoking, this signature was found only in cancer types in which tobacco smoking increases risk and mainly in those derived from epithelia directly exposed to tobacco smoke^19^; Middle, the pattern of a mutational signature of lesions from mice exposed to 4NQO, compilation of all 4 samples analyzed; Bottom, the pattern of a mutational signature of lesions from mice exposed to DMBA. The similarity between signature tobacco smoking associated HNSCC and signature 4NQO is 93.9%; and the similarity between signature tobacco smoking associated HNSCC and signature DMBA is only 39.7%. **(c)** Percentage of somatic substitutions located in translated or untranslated in tobacco smoking associated HNSCC patients (left), 4NQO derived lesions (middle) and DMBA derived lesions (right). **(d)** Graphical matrix representation of the individual mutations in 4 syngeneic cell lines (4MOSCs) isolated from lesions from mice exposed to 4NQO. Listed are alterations frequently observed in human HNSCC, and their corresponding percentage of mutations. Mutations (red), or no mutations (blue) are listed in rows and four different cell lines are in column. (**e**) Mutational Plot of *TP53* mutations in 243 HPV-negative tumor samples from TCGA (top) and of 4 syngeneic cell lines (4MOSCs) (bottom). Frequency of mutation is depicted by height of lollipop, while red circle depicts mutations in common between human and mouse HNSCCs.

### Orthotopic oral SCC lesions reflect the complexity of the human HNSCC tumor immune microenvironment (TIME), including exhausted-CD8 T cell infiltration

Transplantation of the 4MOSC cells orthotopically into the tongue of immunocompetent C57Bl/6 mice led to the formation of well-differentiated HNSCC tumors in two of the cell lines, 4MOSC1 and 4MOSC2, which exhibit typical HNSCC histology, as indicated by hematoxylin and eosin (H&E) stained sections and fluorescence cytokeratin 5 staining (**Fig. 2a** and **Supplementary Fig. 2a**). 4MOSC3 and 4MOSC4 cells also formed tumors, but they regressed spontaneously after 2 weeks. This is likely due to their rejection by the host immune system, as judged by their continuous growth upon CD8 T cell depletion (not shown). Thus, we focused our studies on 4MOSC1 and 4MOSC2, with emphasis on investigating whether they have distinct biological properties reflecting human HNSCC. In this regard, since HNSCC has a high propensity to metastasize to locoregional lymph nodes (reviewed in^28^), leading to poor prognosis, we next addressed the metastatic potential of our model. Histological evaluation in H&E stained sections revealed growth of cancer cells in the lymph nodes of mice bearing 4MOSC2 but not 4MOSC1 tumors (**Fig. 2b**). 4MOSC2 tumors also exhibited much higher density of lymphatic vessels staining positive for LYVE-1 than in 4MOSC1 (**Fig. 2c**), which is aligned with the strong correlation between intratumoral lymphangiogenesis and metastasis in human HNSCC (reviewed in^29^).

**Fig. 2.**
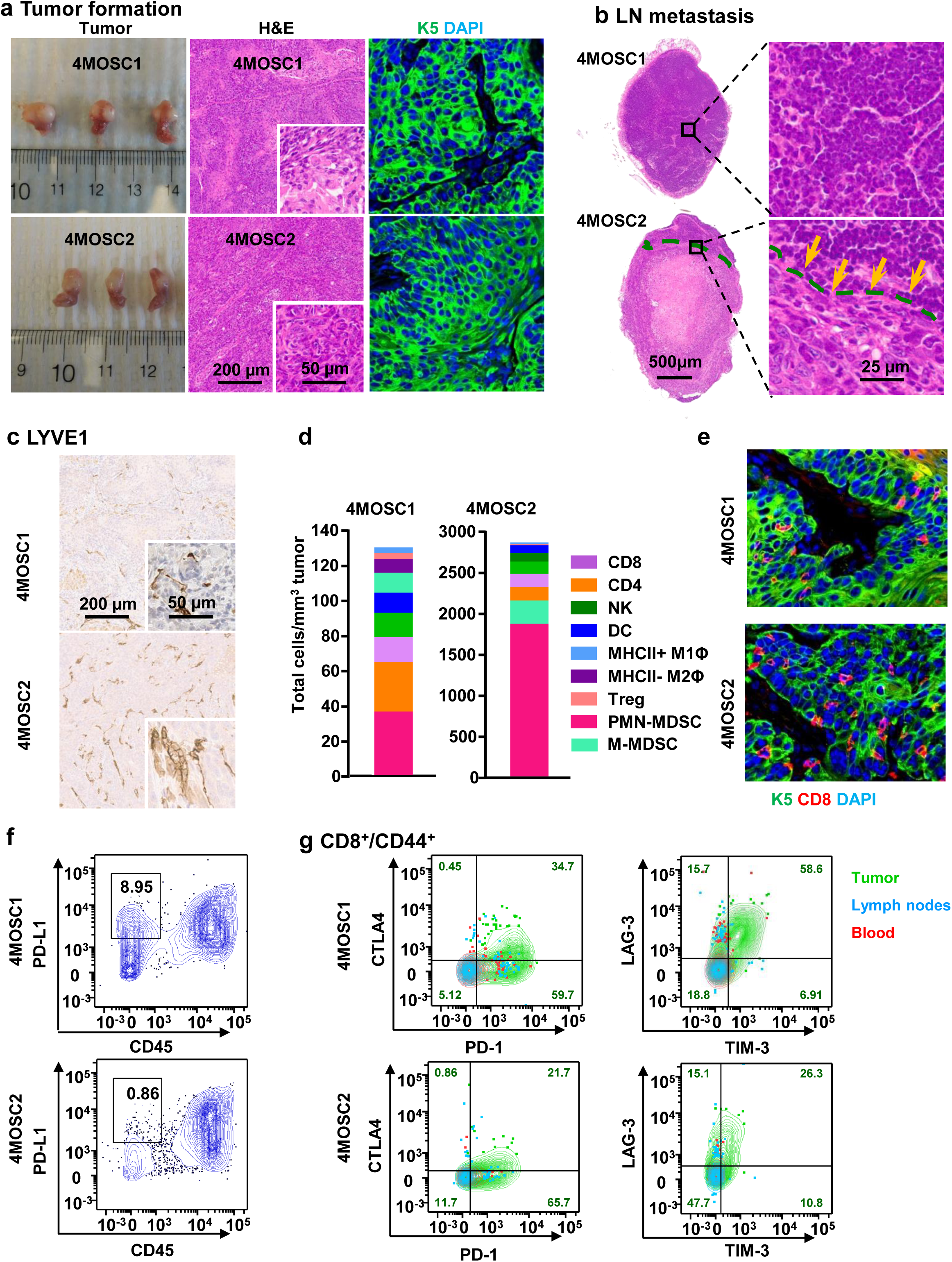
Characterization of 4NQO-induced murine oral squamous cell model. **(a)** Left panel, C57Bl/6 mice were implanted with 1×10^6^ of either 4MOSC1 or 4MOSC2 cells into the tongue. Tongue lesions when the tumor volume reached approximately 100 mm^3^. Middle panel, representative H&E-staining of histological tissue sections from mouse tongues with 4MOSC1 or 4MOSC2 tumors. Right, representative pictures of tumors stained to show expression of cytokeratin 5 (CK5, green) and DAPI (blue). **(b)** Top panel, representative H&E stain of a non-metastatic cervical lymph node from mice with 4MOSC1 tumors. Bottom panel, representative H&E stain of a metastatic cervical lymph node from mice with 4MOSC2 tumors. Metastatic growth of 4MOSC2 cells into the lymph node is depicted with a dotted line in the bottom area. **(c)** Representative tumor tissue sections (right) stained for LYVE1 by immunohistochemistry in 4MOSC1 or 4MOSC2 tumors. **(d)** 4MOSC1 or 4MOSC2 tumors were isolated from mice and mechanically and enzymatically digested. Single cell suspensions were stained with fluorescent labeled antibodies for flow cytometry analysis of each immune cell population. Absolute number of immune cells was quantified using BD Trucount Absolute Counting Tubes. Shown is the average number of live cells infiltrating per mm^3^ of tumor (n = 3). **(e)** Immunofluorescent staining of CK5 and CD8 to show squamous cell character of the lesion and CD8 infiltration in mice with 4MOSC1 or 4MOSC2 tumors, respectively (n = 3) (CK5, green; CD8, red; DAPI, blue). **(f-g)** 4MOSC1 or 4MOSC2 tumors were isolated from mice and mechanically and enzymatically digested. Single cell suspension was then stained with CD45, CD3, CD8, CD44, PD-L1, PD-1, CTLA-4, LAG-3 and TIM-3 fluorescent labeled antibodies and analyzed by flow cytometry. Shown are representative flow cytometry plots of (**f**) the frequency of tumor cells (CD45 negative) expressing PD-L1 and (**g**) the frequency of CD8^+^/CD44^+^ cells expressing inhibitory receptors PD-1, CTLA-4, LAG-3 and TIM-3 (n = 4). Contour plots of lymphocytes from tumor (green), and corresponding cervical lymph nodes (blue), and blood (red) are overlaid and the frequencies of CD8^+^/CD44^+^ T cells expressing each inhibitory receptor are shown.

By flow cytometry analysis, we found that the immune cells infiltrating the TIME comprises of cytotoxic T cells (CD8), helper T cells (CD4), regulatory T cells (Treg), natural killer cells (NK), macrophages (M1Φ and M2Φ), as well as polymorphonuclear myeloid-derived suppressor cells (PMN-MDSC), and monocytic myeloid-derived suppressor cells (M-MDSC)^30^ (**Fig. 2d**). Notably, although the ratios of immune infiltration were similar, 4MOSC2 tumors had a considerably higher level of infiltration than 4MOSC1 (**Fig. 2d**). Though cytotoxic CD8 T cells infiltrate both tumors at similar proportions relative to other immune cells, immunofluorescence staining showed more abundant distribution within 4MOSC2 tumor cells (**Fig. 2e**). To determine whether this immune infiltration is associated with antigen-driven immunogenicity, we next investigated whether the 4MOSC tumors could generate memory immune responses. Initial exposure of mice to tumor cell antigens was achieved by first irradiating tumor cells and then injecting into the tongue of C57Bl/6 mice with or without polyinosinic-polycytidylic acid (poly IC) as an immune adjuvant. Irradiated 4MOSC1 and 4MOSC2 cells did not form tumors, and mice vaccinated with irradiated tumor cells alone or irradiated tumor cells with poly IC failed to form tumors when they were subsequently re-challenged with non-irradiated cancer cells, while naïve mice and mice with poly IC alone still formed tumors. This suggests that the mice were able to develop an immunological memory to 4MOSC antigens even in the absence of an immune adjuvant, indicating that these syngeneic HNSCC cell lines are highly immunogenic (**Supplementary Fig. 2b-e**).

These findings indicate that these mice are capable of generating adaptive immune responses against 4MOSC tumor antigens. However, 4MOSC tumors still grow and lead mice to succumb to disease, implying that these tumors can evade immunity by inducing an immune suppressive microenvironment. The expression of programmed death ligand 1 (PD-L1), the ligand for the T-cell inhibitory receptor PD-1, is often high in HNSCC patients (46-100% of tumors)^31^ and has been shown to suppress cytotoxic T cells that destroy tumors and also serves as a biomarker (reviewed in^32^) predicting a better response to anti-PD-1 therapy. We found that PD-L1 is constitutively expressed on both tumor (CD45^−^) and immune cells (CD45^+^), but the frequency of 4MOSC1 tumor cells that expressed PD-L1 was much higher than the frequency of 4MOSC2 cells expressing PD-L1 (**Fig. 2f**). Activated (CD44^+^) CD8 T cells infiltrating both 4MOSC1 and 4MOSC2 tumors exhibited characteristic T-cell exhaustion markers, PD-1, cytotoxic T lymphocyte associated protein 4 (CTLA-4), T-cell immunoglobulin mucin 3 (TIM-3), and lymphocyte activation gene 3 (LAG-3) (**Fig. 2g**). Interestingly, there were higher T-cell exhaustion markers in tumor-infiltrating lymphocytes (TILs, green) than in lymph nodes (blue) or blood (red) in both tumors (**Fig. 2g**).

### Most syngeneic HNSCC tumors are refractory to anti-PD-1 immune checkpoint blockade

To interrogate whether blocking the interaction of PD-1 and PD-L1 could cause tumor regression, we first studied 4MOSC1 tumors, the syngeneic HNSCC cell line that has higher PD-L1 expression. Most mice showed an initial decreased tumor volume after anti-PD-1 treatment (**Fig. 3a**), with a consequent increased overall survival (**Fig. 3b**) (p<.001). Interestingly, however, 75% of mice that initially responded to anti-PD-1 (partial response, PR) showed tumor relapse and eventually succumbed to disease burden, while 10-20% of the mice showed complete responses (CR) (**Fig. 3a-b,** and **Supplemental Fig. 3a**). 4MOSC1 tumors from mice treated with anti-PD-1 showed significantly higher CD8 infiltration (p<0.01) compared to tumors from untreated mice by FACS analysis of tumor infiltrating leukocytes (TILs) and by immune fluorescence analysis of treated tissues (**Fig. 3c** and **3d**, respectively). All responses to anti-PD-1 were abolished if CD8 T cells were eliminated from mice (**Fig. 3e** and **Supplemental Fig. 3b**). Together, these data indicate a CD8-dependent anti-PD-1 response in mice with 4MOSC1 tumors, but with limited durable disease control or tumor regression, which is similar to the clinical response to anti-PD-1 therapies in HNSCC patients^14,16^. To our surprise, although metastasis was not observed in 4MOSC1 control mice (see above), the cervical lymph nodes of CD8-depleted mice with 4MOSC1 tumors showed tumor invasion, as indicated by cytokeratin 5 staining (**Fig. 3f**) and visualization with H&E-staining (**Supplemental Fig. 3c**) suggesting that immune surveillance may prevent the metastatic spread of this tumor. In contrast, mice bearing 4MOSC2 tumors failed to respond to anti-PD-1 treatment (**Fig. 3g** and **Supplemental Fig. 3d**), suggesting 4MOSC2 serves as a less differentiated, high immune cell-infiltrated, metastatic, and PD-L1-low, and anti-PD-1 resistant model (**Fig. 3g**, and **Supplemental Fig. 3d**).

**Fig. 3.**
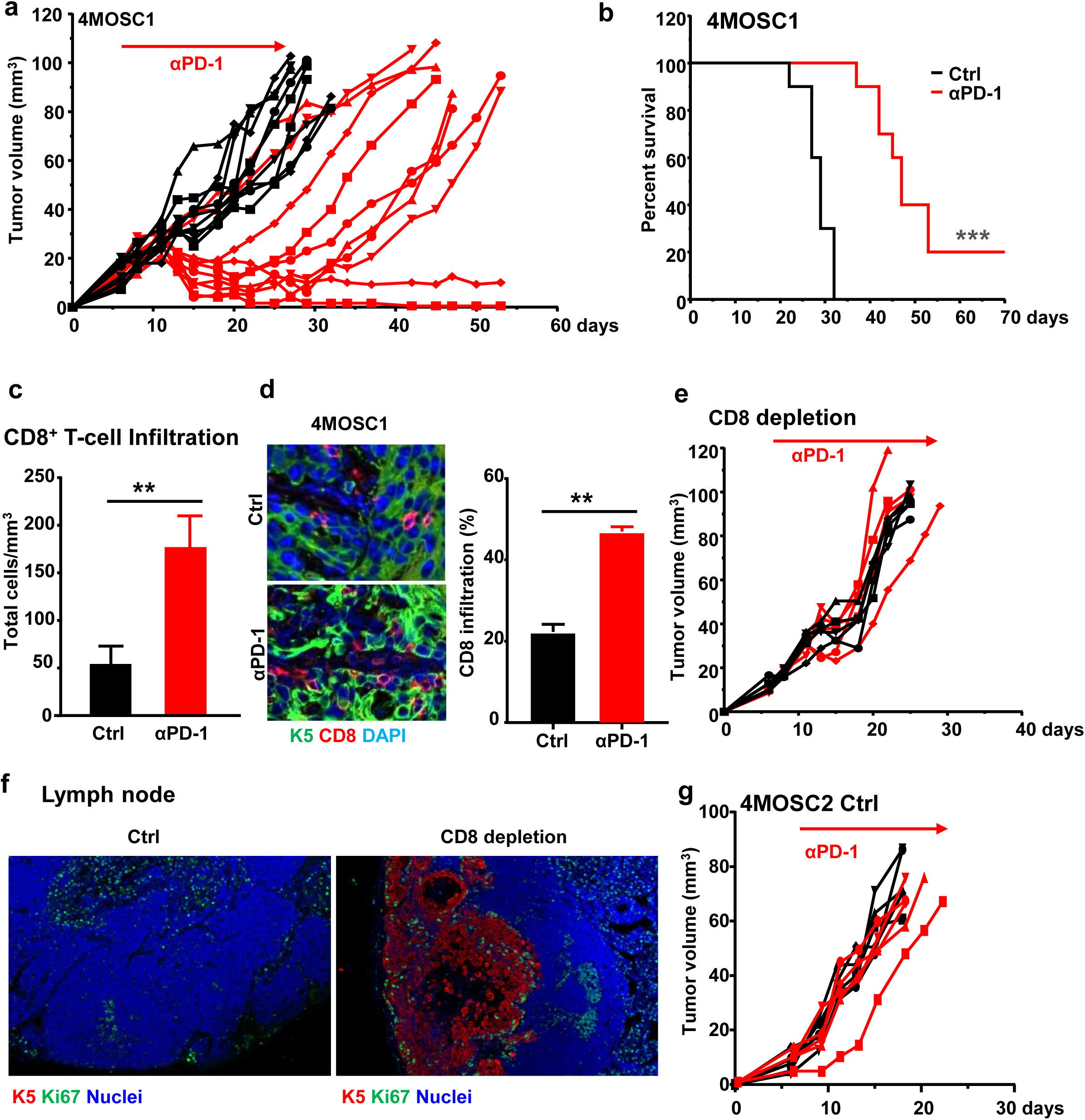
Variable responses to anti-PD-1 in mice with 4MOSC1 tumors. **(a)** Anti-tumor efficacy of anti-PD-1 in mice with 4MOSC1 tumors. C57Bl/6 mice were implanted with 1×10^6^ of 4MOSC1 cells into the tongue. After the tumors reached ∼30 mm^3^, mice were treated IP with 10mg/kg of isotype control (black) or 10mg/kg of anti-PD-1 (red) (n = 10 per group). Individual growth curves of 4MOSC1 tumor-bearing mice are shown. **(b)** A Kaplan-Meier curve showing the effect of anti-PD-1 on the survival of mice with 4MOSC1 tumors. The death of animals occurred either naturally, when the tumor compromised the animal welfare, or when the tumor volume reached 100 mm^3^ (n = 10, ***p < 0.001, Log-Rank/Mantel-Cox test). **(c)** 4MOSC1 tumors from control or anti-PD-1-treated mice were isolated and mechanically and enzymatically digested. Single cell suspensions were stained with CD45, CD3, and CD8 fluorescent labeled antibodies for flow cytometry analysis of CD8 T cells. Absolute number of immune cells was quantified using BD Trucount Absolute Counting Tubes. Shown is the average of the number of live CD8 T cells infiltrating per mm^3^ of tumor (n = 3, **p < 0.01, Student’s t-test). **(d)** Immunofluorescent staining of CD8 highlights an increase in CD8 T cell recruitment with anti-PD-1 treatment (n = 3, p<0.01; K5, green; CD8; red, DAPI; blue). **(e)** Dependency of anti-PD-1 on CD8 T cells. C57Bl/6 mice were treated with CD8 T cell depleting antibody daily for 3 days before tumor implantation and then once a week after. Mice were then implanted with 1×10^6^ of 4MOSC1 cells into the tongue. After the tumors reached ∼30 mm^3^, mice were treated IP with 10mg/kg isotype control (black) or 10mg/kg anti-PD-1 (red) (n= 5 per group). Individual growth curves of 4MOSC1 tumor-bearing mice are shown. **(f)** Immunofluorescence staining of CK5 and Ki67 in cervical lymph nodes of control or CD8-depleted 4MOSC1-bearing mice was used to determine whether tumor cells metastasized. Metastatic lesions in the lymph nodes also showed abundant Ki-67^+^ proliferating tumor cells. **(g)** C57Bl/6 mice were implanted with 1×10^6^ of 4MOSC2 cells into the tongue. After the tumors reached ∼30 mm^3^, mice were treated IP with 10mg/kg isotype control (black) or 10mg/kg anti-PD-1 (red) (n = 5 per group). Individual growth curves of 4MOSC2 tumor-bearing mice are shown.

### Immune regulatory activity of intratumoral (*in situ*) delivery of ICB antibodies

A defining feature of most HNSCCs is the superficial and mucosal localization of the disease. Unlike many other cancer types, most HNSCC patients have tumors that can be readily visualized and accessed by surgeons, providing an opportunity to use intratumoral (IT) drug delivery. To investigate whether IT injection of anti-PD-1 has improves activity in our model, we compared the effectiveness of using a lower dose anti-PD-1 treatment with standard systemic delivery. We found that mice treated with just half the dose of anti-PD-1 locally showed similar anti-tumor responses compared to mice with full dose systemic treatment (**Fig. 4a**). Furthermore, immunofluorescence analysis revealed that IT delivery led to significantly higher PD-1 antibody distribution in tumors and cervical lymph nodes and lower distribution in the spleen as a peripheral organ when compared to systemic delivery (**Fig. 4b**).

**Fig. 4.**
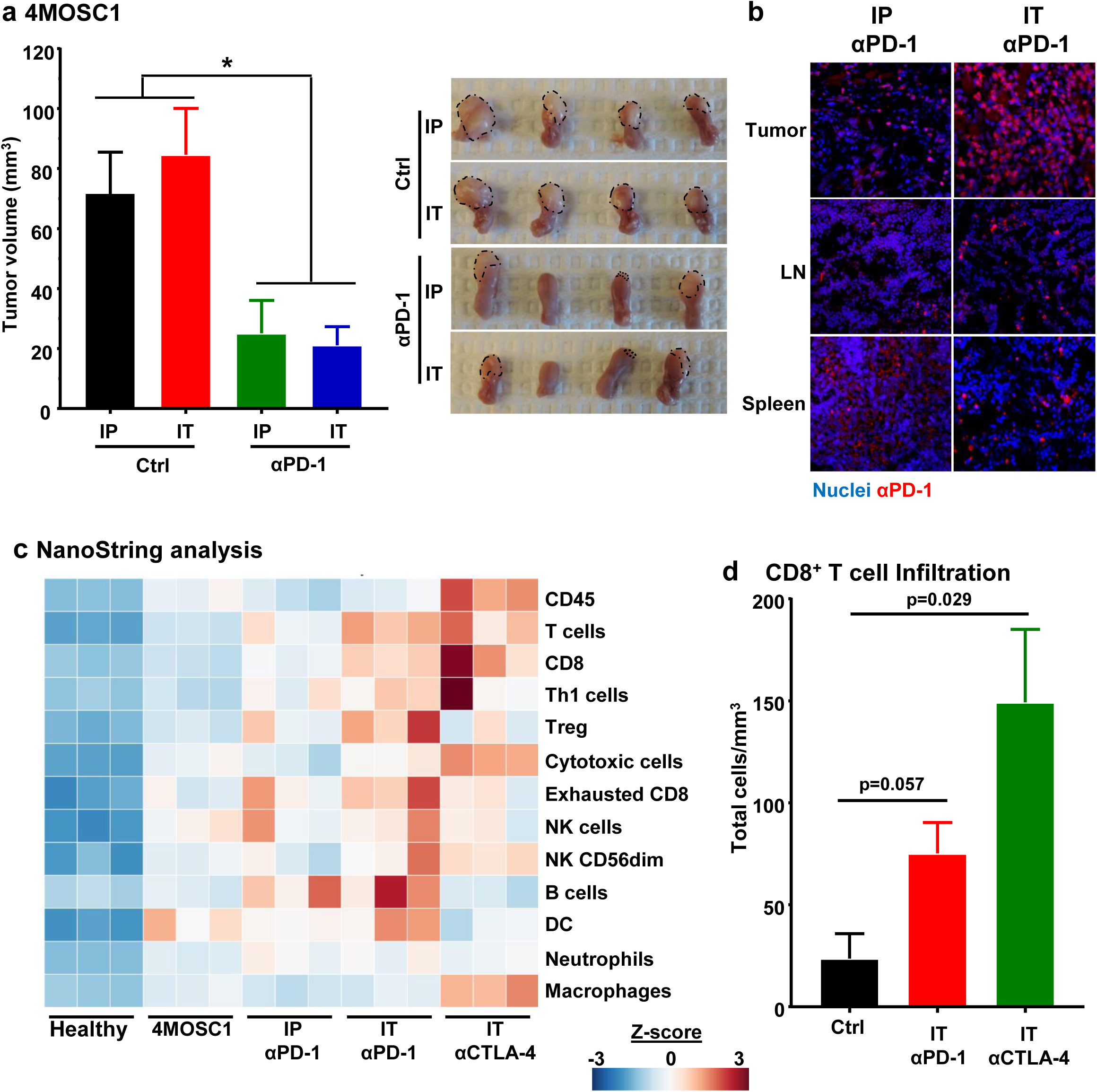
Efficacy of intratumoral delivery of immune oncology agents. **(a)** Left panel, C57Bl/6 mice were implanted with 1×10^6^ of 4MOSC1 cells into the tongue. After the tumors reached ∼30 mm^3^, mice were either treated IP with PBS (black) or by intratumoral (IT) delivery of PBS (red), IP with 10mg/kg anti-PD-1 (green) or IT with 5mg/kg anti-PD-1 (blue) (n = 5 per group). The tumor growth curves were compared by the longitudinal data analysis method (two-sided). Shown is the average volume of each tumor with error bars representing standard error (* p<0.05, when compared with the treatment control group; n = 5 per group). Right panel, representative pictures of mouse tongues from mice in panel **A** with tumors depicted with a dotted line. **(b)** Distribution of anti-PD-1 antibody in mice with 4MOSC1 tumors using IP or IT delivery of the treatment. Staining for anti-hamster IgG showed the localization of anti-PD-1 antibody in the tongue, lymph nodes and spleen of treated mice. **(c)** RNA from each tumor was isolated and comprehensive immune profiling was analyzed using the NanoString nCounter PanCancer Mouse Immune Profiling gene expression platform. The Advanced Analysis module of the nSolver software was used to analyze genes associated with listed immune cells and given a score. Shown is the Z-score of each cell profile score (n = 3 mice per group). **(d)** 4MOSC1 tumors were isolated from mice treated with IT anti-PD-1 or anti-CTLA-4 and mechanically and enzymatically digested. Single cell suspensions were stained extracellularly with CD45, CD8, and CD8 fluorescent labeled antibodies and analyzed by flow cytometry (n = 5). The frequency of CD8+ cells was quantified following treatment with anti-PD-1 or anti-CTLA-4 (n = 5, p value indicated, Student’s t test).

To investigate if there are local, differential immune signature alterations associated with IT drug delivery, we performed a comprehensive immune profiling using the nCounter PanCancer Mouse Immune Profiling gene expression platform (NanoString Technologies). Using 770 immune-related genes, we profiled immune cells infiltrating the tumor following treatment with systemic or intratumoral anti-PD-1, and compared it with another FDA-approved immunotherapy, CTLA-4 blockade. Relative to the tongues from healthy mice, mice bearing 4MOSC1 tumors have elevated expression of a majority of immune cell-associated genes. While systemic anti-PD-1 treatment increased the T cell signature, IT delivery enhanced it further while also increasing gene expression of natural killer cells and genes related to cytotoxic immune cells (**Fig. 4c**). Interestingly, albeit used initially as a control, this analysis revealed that IT treatment with anti-CTLA-4 led to even more robust T cell, cytotoxic cell, and macrophage responses. In addition, while anti-PD-1 increased Treg associated gene signatures, anti-CTLA-4 appears to diminish it (**Fig. 4c-d**).

### Complete response to anti-CTLA-4 ICB in the majority of 4MOSC1 syngeneic HNSCC lesions

Given that immune stimulatory effects were enhanced following anti-CTLA-4 treatment compared to anti-PD-1 treatment, we sought to determine whether mice with 4MOSC1 tumors can also respond to CTLA-4 blockade. CTLA-4 blockade systemically and IT elicited a robust anti-tumor effect, with 90% of the mice exhibiting a CR (**Fig. 5a** and **Supplemental Fig. 4a**) and efficiently resisted engraftment when re-challenged with fresh 4MOSC1 cells (data not shown). Similar to anti-PD-1 treatment, IT delivery resulted in significantly higher anti-CTLA-4 antibody distribution in tumors and cervical lymph nodes and lower distribution in the spleen (**Fig. 5b**). Anti-tumor immunity of anti-CTLA-4 is also CD8 dependent, as CTLA-4 inhibition resulted in significantly increased infiltration of CD8^+^ T cells (**Fig. 5c**), and its anti-tumor activity was abolished by depletion of CD8 T cells (**Supplemental Fig. 4b**). Additionally, flow cytometry analysis of the tumor revealed that following treatment with anti-CTLA-4, there is a significant decrease in immunosuppressive regulatory FoxP3^+^ T cells (Tregs) (**Figs. 5c** and **5d**). In contrast to the effect of anti-CTLA-4, anti-PD-1 treatment led to significant increases in Tregs and FoxP3 expression in the TILs. Of interest, 4MOSC2 tumors also failed to respond to anti-CTLA-4 treatment (**Supplemental Fig. 4c**), providing a model that is resistant to both forms of immunotherapy for future exploration of immunotherapy resistance and the use of strategic combinatorial modalities.

**Fig. 5.**
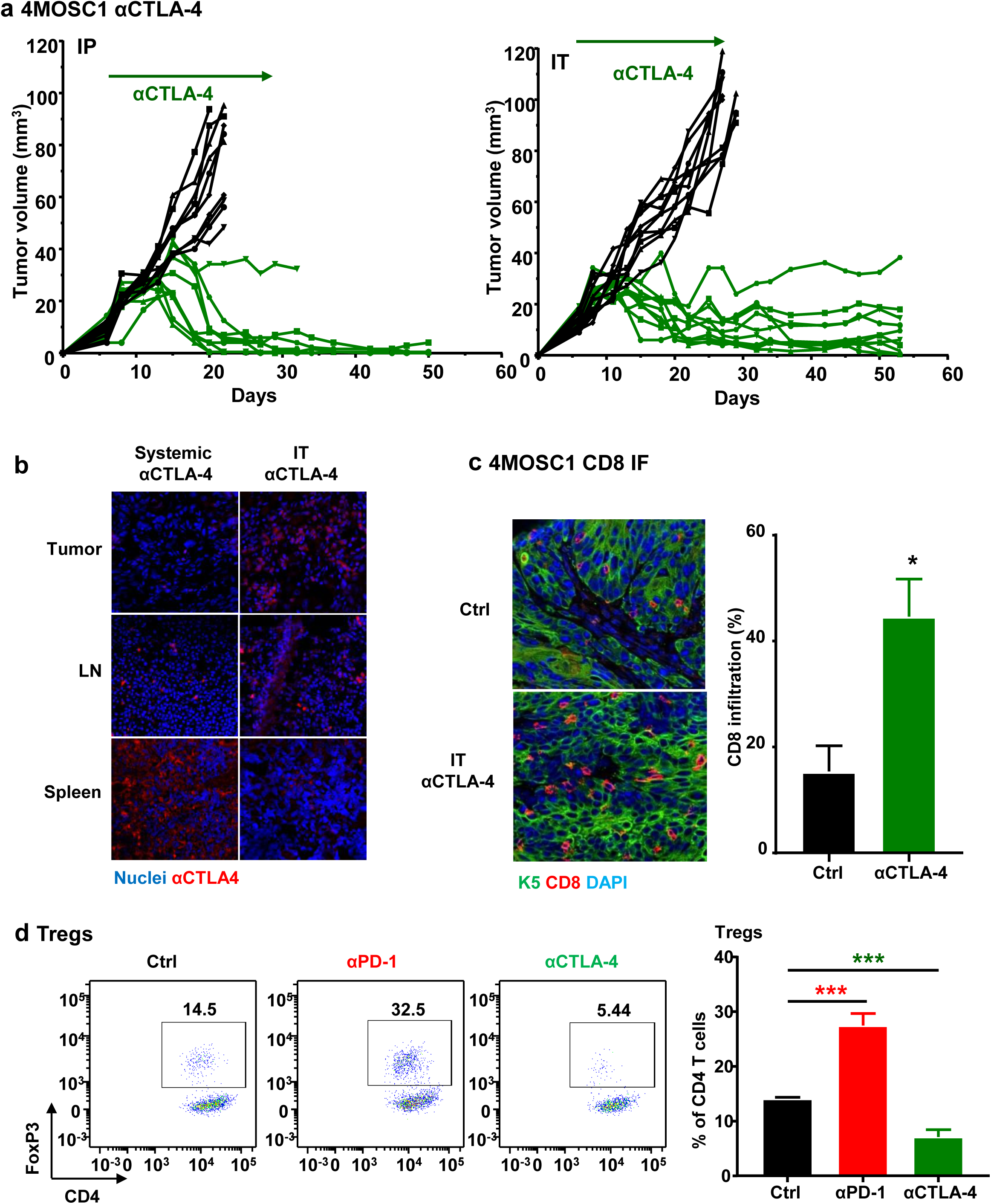
Mice with 4MOSC1 tumors show nearly complete response to anti-CTLA-4. **(a)** Anti-tumor efficacy of anti-CTLA-4 for mice with 4MOSC1 tumors. C57Bl/6 mice were implanted with 1×10^6^ of 4MOSC1 cells into the tongue. After the tumors reached ∼30 mm^3^, mice were treated 10mg/kg of isotype control (black) or anti-CTLA-4 (green) (n = 10 per group) for IP administration (left), and 5mg/kg of isotype control (black) or anti-CTLA-4 (green) (n = 10 per group) for IT administration (right). Individual growth curves of 4MOSC1 tumor-bearing mice plotting primary tumor growth are shown. **(b)** Shown is the immunofluorescent staining of the distribution of anti-CTLA-4 antibody for mice with 4MOSC1 tumors using IP or IT delivery of the treatment. Staining for anti-hamster IgG (red) showed the localization of anti-CTLA-4 antibody in the tongue, lymph nodes and spleen of treated mice. DAPI staining for nuclei is shown in blue. **(c)** Quantification of CD8 T cells with or without anti-CTLA-4 treatment was done by immunofluorescent staining of tumor (CK5) in the tongue. CD8 positivity was quantified by Qupath software counting the number of cells showing CD8 staining in at least 3 regions of interest (ROI) for each condition (n = 3, *p<0.05). **(d)** 4MOSC1 tumors from mice treated with or without anti-PD-1 and anti-CTLA-4 were isolated and mechanically and enzymatically digested. Single cell suspensions were stained extracellularly with CD45, CD3, CD4 and intracellularly with FoxP3 fluorescent labeled antibodies and analyzed by flow cytometry (n = 5). Left panel, a representative flow cytometry plot from one mouse showing the frequency of Tregs (CD4^+^FoxP3^+^) out of CD4^+^ cells is shown. Right panel, the frequency of Tregs out of CD4^+^ cells was quantified following treatment with anti-PD-1 or anti-CTLA-4 (n = 5, ***p<0.001, Student’s t test).

## Discussion

HNSCC is an immunosuppressive disease, in which the tumor deploys multiple mechanisms to evade immune surveillance and antitumor immune responses through the accumulation of immunosuppressive cytokines, impairment of cytotoxic activity and antigen-presenting function, and induction of T cell exhaustion (reviewed in^12^). Based on this knowledge, numerous immunotherapeutic strategies were developed, including ICB, cancer vaccines, therapeutic cytokines, adoptive T-cell transfer, and adjuvants that may trigger innate immune responses such as TLR and STING agonists (reviewed in^12^). Clearly, suitable experimental systems that can model clinical responses are urgently needed to study and improve the effectiveness of immune oncology approaches in HNSCC. Here, we developed a panel of C57Bl/6-derived syngeneic cells that resemble human HNSCCs closely with unique features: (i) The HNSCC cells have nearly identical tobacco-associated mutational signatures and genomic aberrations; (ii) they can be orthotopically transplanted into the tongue of immunocompetent C57Bl/6 mice; (iii) the tumors are histologically HNSCCs with abundant lymphangiogenesis and potential for lymph node metastasis; and (iv) the tumors exhibit abundant immune infiltration and are immunogenic, the latter as judged by their ability to induce immunological memory when used to vaccinate mice. These novel animal models may provide an opportunity to investigate the mechanisms driving intrinsic and acquired resistance to IO agents, as well as to identify novel therapeutic options increasing the response of currently available immunotherapies in HNSCC patients.

Tobacco use is one of the major risk factors for initiation and progression of HNSCC, and serves as an important prognostic factor for survival and mortality after cancer diagnosis (reviewed in ^33^). In a recent study, we analyzed somatic mutations and DNA methylation in 5243 samples comprising of cancers for which tobacco smoking confers an elevated risk, which helped define the human tobacco-associated cancer signature^19^. Remarkably, although tobacco smoke is made up of thousands of chemicals, including more than 60 carcinogens^17^, we found that 4NQO exposure was sufficient to mimic the tobacco carcinogenic signature. This is supported by compelling evidence demonstrating that 4NQO-induced SCC lesions exhibit near identical association (∼94%) with the tobacco mutational landscapes, recapitulating human HNSCC. This is in contrast with the mutational signature caused by DMBA (<40% similarity), which although representing a widely used tobacco carcinogen^26^, may not be as effective as 4NQO in reflecting the human tobacco-associated genetic signatures^19^.

In this regard, currently available syngeneic HNSCC models include SCCVII cells, HPV+ SCC cells designated MEER^34^ and a panel of mouse HNSCC cell lines from DMBA-treated mice (MOC1 and MOC2)^35^. Though widely used, SCCVII cells are in fact derived from a spontaneously formed skin SCC lesion in C3H mice^36^. MEER and MOC1/MOC2 models develop tumors in immune competent C57Bl/6 mice when implanted in the flanks, which may not reflect the HNSCC TIME. In addition, these tumors are driven by *Ras* oncogenes (*Kras* in MEER and MOC2, and *Hras* in MOC1), which are potent oncogenic drivers but infrequently mutated in human HNSCC^34,35,37^, and therefore likely not reflecting the tobacco-induced carcinogenic process driving human HNSCC. Taken together, the unique features of our novel syngeneic HNSCC animal model provide a resource to investigate novel IO pre-clinical approaches for HNSCC treatment.

Seminal studies have shed light on T cell exhaustion in human cancers, where CD8 T cells lose proliferative capacity, the ability to produce tumor necrosis factor (TNFα), interleukin-2 (IL-2), and interferon-γ (IFNγ), and upregulation of inhibitory checkpoint receptors, such as PD-1 and CTLA-4^9-11^. Recently, the successes of ICB to reverse T cell exhaustion in multiple cancers illustrates the potential of therapeutic strategies targeting these negative regulatory pathways^38^. In the clinic, PD-1 blockade offers 10∼20% clinical improvement in HNSCC^13-15^, which was modeled similarly in our study where anti-PD-1 led to regression of 4MOSC1 tumors in only ≤20% of mice. The increase in CD8 T cells seemed to provide only temporary cytotoxic activity in mice treated with anti-PD-1, as we observed reoccurrence of tumors in the majority of treated mice. Surprisingly, we saw enhanced anti-tumor responses with anti-CTLA-4 treatment, where most 4MOSC1 tumor bearing mice showed complete responses and no tumor reoccurrence. The resulting increase in CD8 T cells following anti-CTLA-4 treatment confirmed that targeting checkpoints may revitalize immunological effect of exhausted T cells, at least at the cellular level. One explanation for these strikingly different responses could be that PD-1 blockade may induce compensatory upregulation of FoxP3^+^ Treg cells^39,40^, as it occurred in our anti-PD-1-treated mice but not in anti-CTLA-4-treated mice. In fact, CTLA-4 inhibition led to significantly lower levels of FoxP3^+^ Treg cells in the tumors. In this regard, while both blocking antibodies can lead to cytotoxic CD8 T cell responses, anti-CTLA-4 may provide additional anti-tumor immunity by depleting Tregs that mediate an immune-suppressive environment (reviewed in^41^). Moreover, PD-1 blockade predominantly activates T cells within the tumor, whereas anti-CTLA-4 may activate T cells primarily in the lymph nodes^41^, in which high levels of anti-CTLA-4 can be achieved by IT delivery. These and yet to be identified mechanisms may underlie the increased response to anti-CTLA-4 in some anti-PD-1 refractory HNSCC lesions, whose elucidation may provide biomarkers for the selection of patients that may benefit from anti-CTLA-4 treatment after failing to anti-PD-1 therapy.

One limitation of using anti-CTLA-4 for HNSCCs in the clinic is its toxicity^42^. Systemic delivery of IO agents have been shown to be responsible for severe immune related adverse events (irAEs), such as colitis, dermatitis, uveitis, and hypophysitis^42^. These adverse events are very toxic, at times irreversible and can even be life-threatening. With this in mind, IT injection may enhance tumor-specific T cell responses while reducing significant systemic exposure to healthy tissue and off-target toxicities^43,44^. In addition, IT immunotherapy usually causes *in situ* priming of antitumor immunity, which may allow a patient’s own tumor cells to be used as a therapeutic vaccine^43,44^. In our study, a lower dose of IT anti-PD-1 showed similar therapeutic effects as systemic delivery of a higher dose, and IT anti-CTLA-4 led to complete regression of most 4MOSC1 tumors that are primarily refractory to anti-PD-1. Additionally, IT injection led to higher distribution of the antibody in the tumor and cervical lymph nodes, but less in the spleen as a surrogate for distribution in peripheral organs. This suggests that the IT route, which is feasible in HNSCC, may serve as a more effective and less toxic therapeutic strategy for this tumor type, a possibility that may have readily applicable clinical implications, and hence warrant further investigation.

Certainly, some HNSCC tumors have minimal immune infiltration, and may require a multipronged approach to facilitate immune recruitment and activation of the anti-tumor immune response^45,46^. Other HNSCC lesions are completely refractory to ICB, even if highly immune infiltrated. In this regard, mice implanted with 4MOSC2 failed to respond to anti-PD-1 and anti-CTLA-4 therapy, likely due to the presence of abundant immune suppressive MDSC^30^ in the TIME, which may restrict DC and/or CD8^+^ T cell function in addition to promoting T cell exhaustion. Therefore, this 4MOSC model system is ideal for investigating mechanisms of immunotherapy resistance, as well as testing novel multimodal immunotherapies and/or optimization of potential combinations of ICB with chemo- and radiotherapies. Altogether, our findings suggest that our novel syngeneic HNSCC animal models, which strongly reflect tobacco-associated HNSCC and typical clinical situations, may provide experimental tools to investigate interplays between HNSCC and the immune system as well as provide unique opportunities to identify more effective therapeutic strategies for tobacco-associated HNSCC, which are associated with poor prognosis and reduced response to most currently available treatment options.

## Materials and Methods

### Reagents

4NQO (4-Nitroquinoline-1-oxide) was purchased from Sigma-Aldrich, dissolved in propylene glycol (Sigma-Aldrich) as a stock solution (4 mg/mL) and stored at 4°C. PD-1 antibody (clone J43, catalog #BE0033-2), CTLA-4 antibody (clone 9H10, catalog #BP0131), isotype antibody (catalog # BE0091) and CD8 depletion antibody (Clone YTS 169.4, catalog #BE0117) were obtained from Bio X Cell (West Lebanon, NH, USA). Fluorochrome-conjugated antibodies were purchased from BioLegend and BD Biosciences.

### Establishment of cell lines and tissue culture

Female C57Bl/6 mice (4–6 weeks of age and weighing 16–18g) were purchased from Charles River Laboratories (Worcester, MA, USA). 4NQO was diluted in the drinking water to a final concentration of 50 μg/mL to animals and was changed weekly. After 16 weeks, all animal cages were reverted to regular water until week 22. Animals were euthanized on week 22 for tissue retrieval. Single lesions were dissected and digested into a single cell suspension according to the protocol in Supplemental Materials.

### DNA sequencing, genomic data analysis, and tobacco signature analysis

Raw sequencing data were aligned to the mm10 reference genome using BWA^47^. Somatic mutations were identified by comparing the sequencing data from each cancer sample to the sequencing data from a normal tissue derived from the tail of one of the mice (all mice were genetically identical). To ensure robustness of the results, a consensus variant calling strategy was leveraged in which somatic mutations were identified using three independent bioinformatics tools: Strelka2^48^, Varscan2^49^, and GATK4 Mutect2^50^. Any mutation found in two out of the three variant callers was considered a bona fide somatic mutation. Additional filtering to remove any residual germline contamination was applied and any mutation found in Mouse Genome Project or shared among all four cancers was discarded. Somatic mutational profiles were derived using the immediate sequencing context by evaluating the base 5’ and the base 3’ to each single point mutation. Additionally, transcriptional strand bias was evaluated by considering all protein coding genes. Mutational signatures were extracted using our previously developed computational framework SigProfiler^24,25^. SigProfiler can be downloaded freely from: https://www.mathworks.com/matlabcentral/fileexchange/38724-sigprofiler.

Gene mutation analyses were performed comparing our 4 syngeneic cells to a HNSCC provisional dataset containing 243 HPV-negative tumor samples from the publicly available consortium, The Cancer Gene Atlas (TCGA)^37^. Mutational plots of p53 mutations observed in characterized HNSCC samples from TCGA and 4 of our syngeneic cell lines were summarized using the ‘lolipop’ mutation diagram generator^51^.

### *In vivo* mouse experiments and analysis

All the animal studies using HNSCC tumor xenografts and oral carcinogenesis studies were approved by the Institutional Animal Care and Use Committee (IACUC) of University of California, San Diego, with protocol ASP #S15195. 4MOSC1 and 4MOSC2 cells were transplanted (1 million per mouse) into the tongue of female C57Bl/6 mice (4–6 weeks of age and weighing 16–18g). When tumors were formed (on day 5-6), the mice were first randomized into groups. For drug treatment, the mice were treated by either intraperitoneal (IP) or intratumoral (IT) injection with isotype control antibody, PD-1 antibody, or CTLA-4 antibody (IP 10mg/kg, IT 5mg/kg, three times a week) for three weeks. The mice were then euthanized after the completion of the treatment (or when control-treated mice succumbed to tumor burdens, as determined by the ASP guidelines) and tumors were dissected for flow cytometric analysis or histologic and immunohistochemical evaluation.

### Immunofluorescence and image quantification

All tissue samples were processed and stained as previously described^52^. Briefly, tissues (tongue, cervical lymph nodes and spleen) were harvested, fixed, and paraffin embedded. Slides were stained for CK5 (Fitzgerald, 20R-CP003) and CD8 (abcam, ab22378) antibodies. Quantification of immune-infiltration was done using QuPath, an open source software for digital pathology image analysis^53^. For the quantification, at least 3 regions of interest (ROI) were selected for each condition and the percentage of positive cells for the CD8 marker was calculated.

### Tumor infiltrating lymphocyte (TIL) isolation and flow cytometry

Tumors were dissected, minced, and re-suspended in complete media (DMEM with 10% FBS and 1% antibiotics) supplemented with Collagenase-D (1mg/mL; Roche) and incubated at 37°C for 30 minutes with shaking to form a single-cell suspension. Tissue suspensions were washed with fresh media and passed through a 100-µm strainer. Samples were washed with PBS and immediately processed for live/dead cell discrimination using BD Horizon™ Fixable Viability Stain 510. Cell surface staining was done for 30 minutes at 4 degrees with the following antibodies (all from BioLegend, San Diego, CA): CD45 (30-F11), CD3 (145-2C11), CD8a (53-6.7), CD4 (RM4-4), NK1.1 (PK136). CD24 (M1/69), MHCII (M5/114.15.2), Ly6-G (1A8), Ly6-C (HK1.4), F4/80 (T45-2342), CD103 (2E7), CD11b (M1/70), CD11c (HL3), PD-1 (29F.1A12), TIM-3 (B8.2C12), and CD44 (IM7). Intracellular staining for inhibitory receptors LAG-3 and CTLA-4 was done using the BD Cytofix/Cytoperm kit and stained with the LAG-3 (C9B7W) and CTLA-4 (UC10-4B9) antibodies. Intracellular staining for FOXP3 was performed using the eBioscience FOXP3/Transcription Factor Buffer Set from Invitrogen and stained with the FOXP3 (MF23) antibody. All flow cytometry data acquisition was done using BD LSRFortessa and analyzed using FlowJo software. TIL count was determined using BD Trucount™ tubes. Immune cells were identified by the following characteristics: cytotoxic T cells (CD45^+^CD3^+^CD8^+^), helper T cells (CD45+CD3+CD4+), Treg (CD45^+^CD3^+^CD4^+^FOXP3^+^), NK cells (CD45^+^CD3^−^NK1.1^+^), macrophages (CD45^+^CD3^−^ NK1.1^−^CD11b^+^CD11c^−^LY6C^low^LY6G^low^CD24^+^F4/80^+^), PMN-MDSCs (CD45^+^CD3^−^NK1.1^−^ CD11b^+^CD11c^−^LY6C^low^LY6G^+^), and M-MDSCs (CD45^+^CD3^−^NK1.1^−^CD11b^+^CD11c^−^ LY6C^+^LY6G^low^).

### NanoString analyses

RNA was isolated from tumor samples using the RNeasy Micro Kit (Qiagen 74004). Hybridization of samples was done according to the NanoString Hybridization Protocol for nCounter XT CodeSet Gene Expression Assays. Samples were run on the nCounter SPRINT Profiler with the nCounter PanCancer Mouse Immune Profiling gene expression platform. Analysis of gene expression was done using the Advanced Analysis module on the nSolver software.

### Statistical Analysis

Statistical data analyses, variation estimation and validation of test assumptions were carried out with GraphPad Prism version 7 statistical analysis program (GraphPad Software, San Diego, CA). All analyses were performed in triplicate or greater and the means obtained were used for independent t-tests, ANOVA, or longitudinal data analysis method. The asterisks denote statistical significance (non-significant or ns, P>0.05; *P<0.05; **P<0.01; and ***P<0.001). All the data are reported as mean ± standard error of the mean (S.E.M.).

## Data Availability

The datasets generated during and/or analyzed during the current study are available from the corresponding author on reasonable request.

## Acknowledgements

This project was supported by grants from National Institute of Dental and Craniofacial Research (NIH/NIDCR, R01DE026644, R01DE026870, and U01DE028227), National Natural Science Foundation of China (81602376, 81520108009, 81621062), and 111 Project of MOE (B14038), China. Mara Gilardi was supported by FIRC-AIRC fellowship for abroad (Italian Foundation for cancer research). Michael Allevato was supported by the Pharmacological Sciences Training Program (5T32GM007752-39 and 5T32GM007752-40). We thank Drs. Trever Greene, Jayanth Shankara Narayanan, Joseph Dolina, Stephen Schoenberger, Zhijun Sun, Dunfang Zhang, Joseph Califano, and Scott M. Lippman for insightful suggestions. We thank the Staff of La Jolla Institute Microscopy Core Facility for professional advice and guidance.

## Author contributions

Z. W., V.H. Wu, M.M. A., and M.G. conducted most experiments described in this study, contributed to the study design, and wrote the corresponding manuscript; Y.H. and L.B.A. conducted the bioinformatics analysis of mutational signatures and gene variants, and contributed to writing the manuscript; J.L.C.-V., L.V.-C., D.M., P.A., J.M. and Y.G. contributed to the isolation and characterization of 4MOSCC cell lines; A.A.M. performed the pathology analysis of all animal studies; A.B.S. and E.E.W.C. contributed to the overall study design and analysis and interpretation of immunological data; Q.C. and J.G.L. contributed to the study design and writing the manuscript, and to the development of methods for the isolation of murine oral cancer cells; J.S.G. provided oversight and direction of the entire project and study design, provided financial support for the study, and wrote the manuscript.

