## supplemental material for "Novel syngeneic animal model of tobacco-associated oral cancer reveals the activity of *in situ* anti-CTLA-4"

**Supplemental Figures and Tables**

**Supplemental Fig. 1. Mutational signatures associated with tobacco smoking.** The somatic mutational profiles of the four 4MOSCs were correlated to (Pearson correlation > 0.93), known mutational signatures in human cancer. The pattern of Signature 4 extracted from cancers associated with tobacco smoking was marked as dark blue columns.

**Supplemental Fig. 2**. **Squamous cell character and immunogenicity of 4MOSC. (a)** Representative pictures of whole tongue tumors stained to show expression of cytokeratin 5 (CK5, green); left, 4MOSC1; right, 4MOSC2. **(b-c)** Memory immune responses induced by vaccination with irradiated 4MOSC cells. 4MOSC1 or 4MOSC2 cells were irradiated with 45 Gy and $1x10^6$ cell were injected into the tongue of C57Bl/6 mice, with (green) or without (blue) polyinosinic-polycytidylic acid (poly IC). Mice injected with non-irradiated 4MOSC cells (black) or mice only treated by poly IC (red) were used as controls. The average tumor volume ± SEM of each group is shown (n = 5 mice per group, ***p<0.001). **(d-e)** Vaccinated mice (green and blue) were re-challenged with $1x10^6$ live 4MOSC cells 6 weeks after. Naïve mice (black) and mice post poly IC treatment (red) were used as controls. The average tumor volume with SEM of each group is shown (n = 5 mice per group, ***p<0.001).

**Supplemental Fig. 3. Histopathological analysis of tongues and cervical lymph nodes from 4MOSC1 or 4MOSC2 tumor-bearing mice. (a-b)** Representative H&E stains of mouse tumors from the experiment in panel 2a and 2e. The H&E stained tissue section of an HNSCC tumor is depicted with a dotted line. **(c)** Top panel, representative H&E stain of a non-metastatic cervical lymph node from mice with 4MOSC1 tumors. Bottom panel, representative H&E stain of a metastatic cervical lymph node from mice with 4MOSC1 tumors after treatment with CD8 T cell-depleting antibody. Metastatic growth of 4MOSC1 cells into the lymph node is depicted with a dotted line in the left area. **(d)** Representative H&E stains of mouse tumors from the experiment in panel 2g. The H&E stained tissue section of an HNSCC tumor is depicted with a dotted line.

**Supplemental Fig. 4. Histological analysis of tongues from 4MOSC1 or 4MOSC2 tumor-bearing mice treated with anti-CTLA-4. (a)** Representative H&E stains of mouse tumors from the experiment in panel 4a. The H&E stained tissue section of an HNSCC tumor is depicted with a dotted line. **(b)** Left panel, anti-CTLA-4 dependency on CD8 T cells. C57Bl/6 mice were treated with a CD8 T cell-depletion antibody, and transplanted with $1x10^6$ 4MOSC1 cells into the tongue. After the tumors reached ~30 $mm^3$, mice were treated IT with 5 mg/kg of isotype control (black) or anti-CTLA-4 (green) (n = 5 per group). Individual growth curves of 4MOSC1 tumor-bearing mice plotting primary tumor growth were recorded. Right panel, representative H&E of mouse tumors from the experiment in left panel. The H&E stained tissue section of an HNSCC tumor is depicted with a dotted line. **(c)** Left panel, antitumor efficacy of anti-CTLA-4 for mice with 4MOSC2 tumors. C57Bl/6 mice were transplanted with $1x10^6$ 4MOSC2 cells into the tongue. After the tumors reached ~30 $mm^3$, mice were treated IT with 5 mg/kg of isotype control (black) or anti-CTLA-4 (green) (n = 10 per group). Individual growth curves of 4MOSC2 tumor-bearing mice plotting primary tumor growth were recorded. Right panel, representative

H&E of mouse tumors from the experiment in left panel. The H&E stained tissue section of an HNSCC tumor is depicted with a dotted line.

**Supplemental Table 1**. **4NQO variant calling results.** Excel file listing variant calling results from DNA sequencing of 4NQO lesions. Column descriptions refer to Genomic Location (Location), Allele, Mutational Consequence, Impact, Symbol, Ensembl Gene, Feature_Type, Transcript Feature, Biotype, cDNA position, CDS_position, Protein position, Amino Acid Mutation, Codon Change, Strand, Transcript Support Level (TSL), Annotation Alternatively Splice Transcripts (APPRIS), and Sorting Intolerant From Tolerant (SIFT). Filter applied to the 4MOSC1-4 samples are used to show only mutations that result in amino acid changes.
