## supplemental figure for "Novel syngeneic animal model of tobacco-associated oral cancer reveals the activity of *in situ* anti-CTLA-4"

#### Supplemental Figure 1:Transcriptional Bias for each individual 4MOSC cell lines

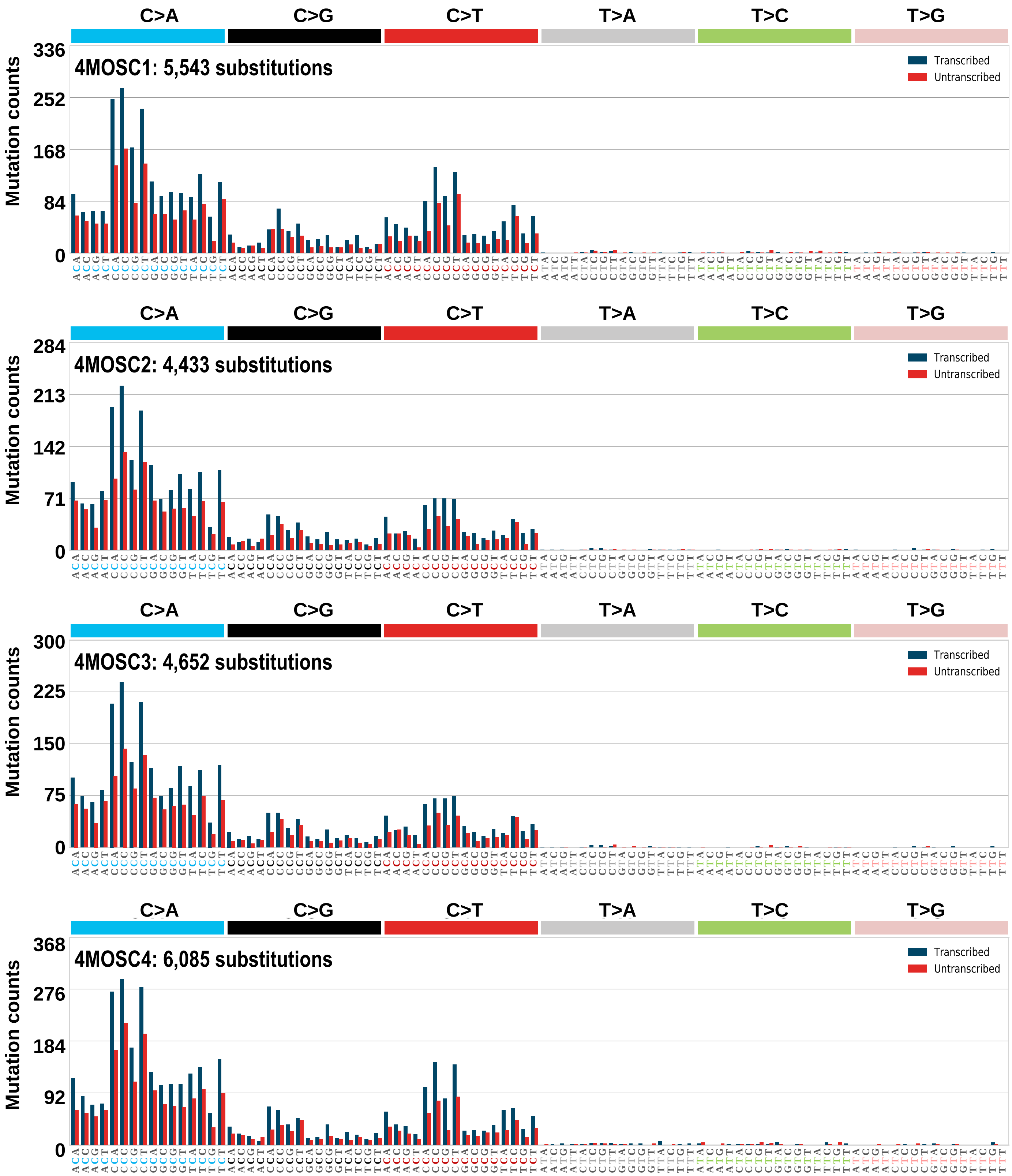

#### Supplemental Figure 2

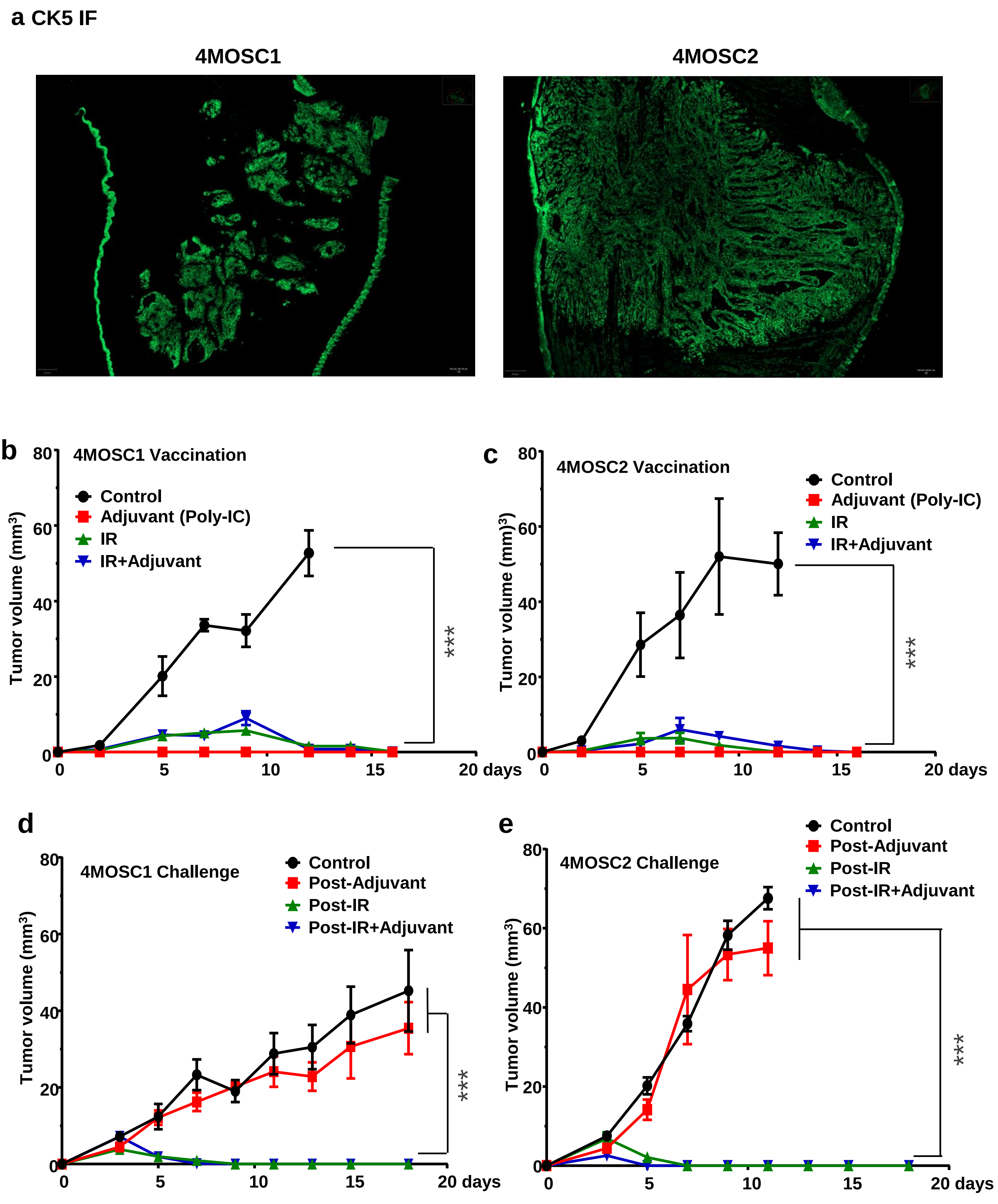

#### Supplemental Figure 3

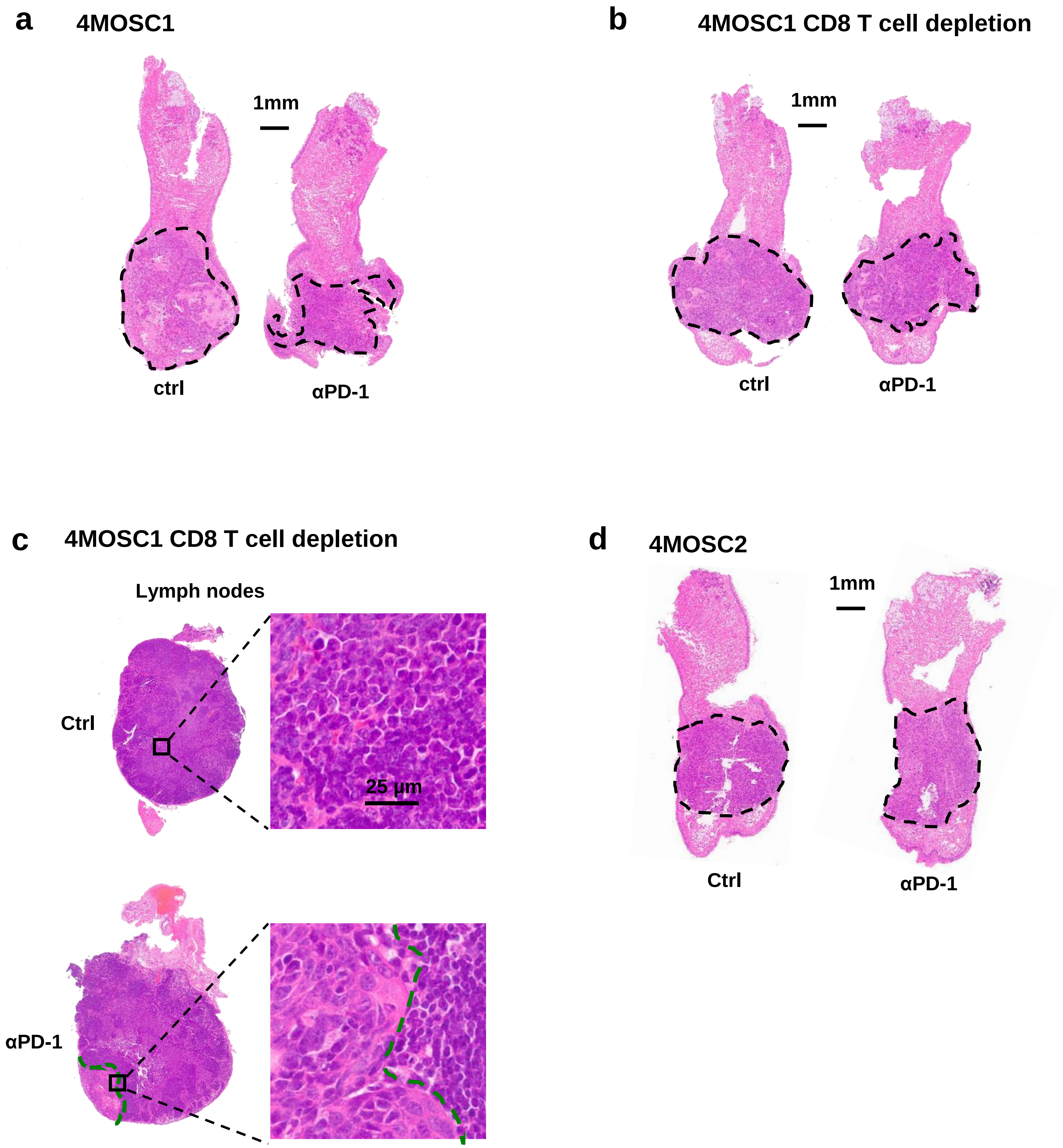

### Supplemental Figure 4

**a** 4MOSC1 αCTLA-4

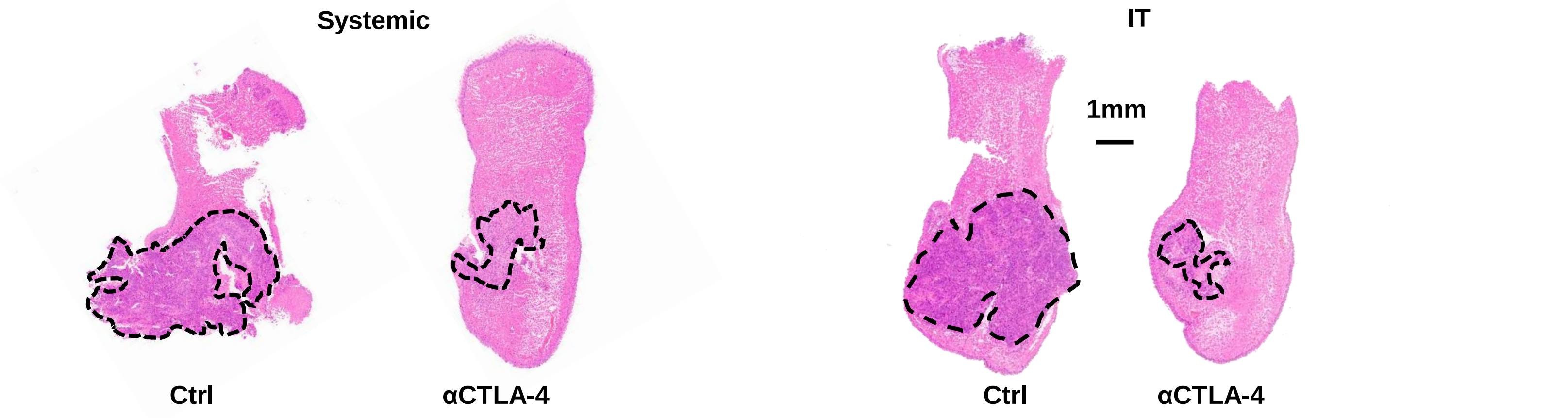


**b** 4MOSC1 CD8 depletion + αCTLA-4

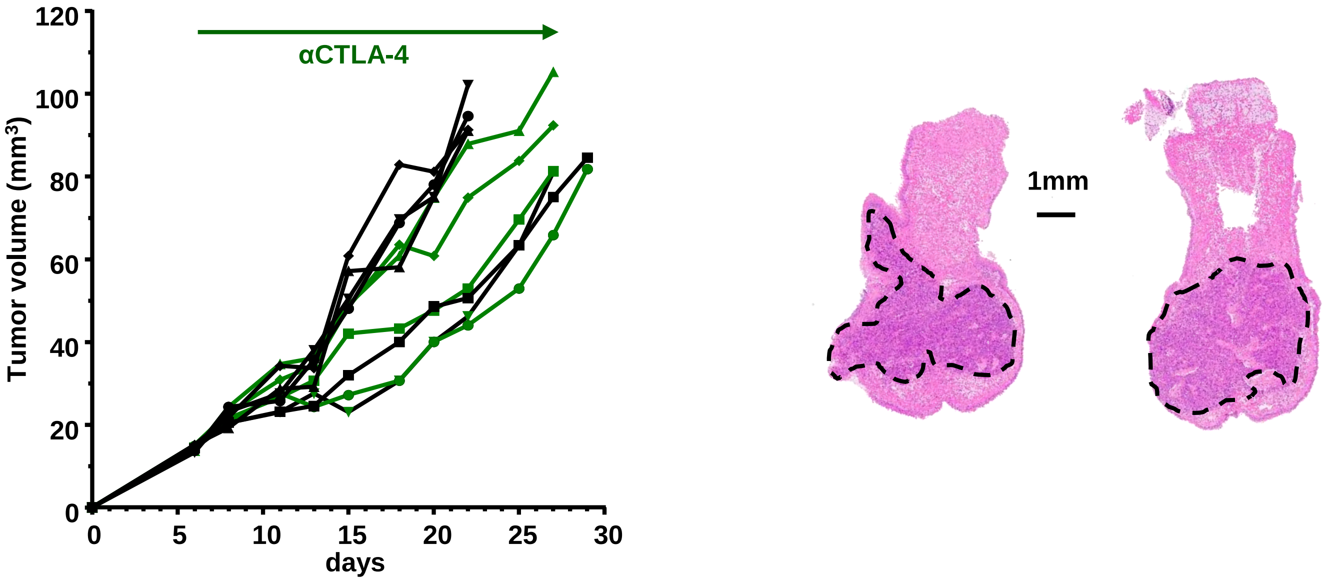


**c** 4MOSC2 αCTLA-4

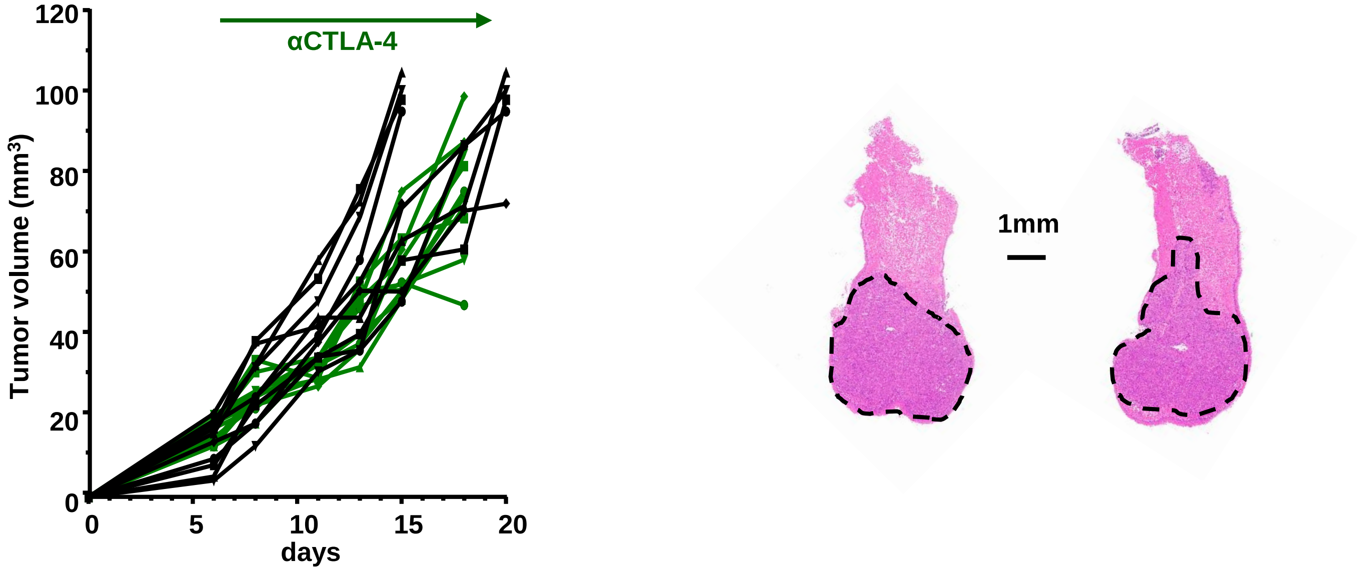
